# Nuclear blebs are associated with destabilized chromatin packing domains

**DOI:** 10.1101/2024.03.28.587095

**Authors:** Emily M. Pujadas Liwag, Nicolas Acosta, Luay Matthew Almassalha, Yuanzhe (Patrick) Su, Ruyi Gong, Masato T. Kanemaki, Andrew D. Stephens, Vadim Backman

## Abstract

Disrupted nuclear shape is associated with multiple pathological processes including premature aging disorders, cancer-relevant chromosomal rearrangements, and DNA damage. Nuclear blebs (i.e., herniations of the nuclear envelope) have been induced by (1) nuclear compression, (2) nuclear migration (e.g., cancer metastasis), (3) actin contraction, (4) lamin mutation or depletion, and (5) heterochromatin enzyme inhibition. Recent work has shown that chromatin transformation is a hallmark of bleb formation, but the transformation of higher-order structures in blebs is not well understood. As higher-order chromatin has been shown to assemble into nanoscopic packing domains, we investigated if (1) packing domain organization is altered within nuclear blebs and (2) if alteration in packing domain structure contributed to bleb formation. Using Dual-Partial Wave Spectroscopic microscopy, we show that chromatin packing domains within blebs are transformed both by B-type lamin depletion and the inhibition of heterochromatin enzymes compared to the nuclear body. Pairing these results with single-molecule localization microscopy of constitutive heterochromatin, we show fragmentation of nanoscopic heterochromatin domains within bleb domains. Overall, these findings indicate that translocation into blebs results in a fragmented higher-order chromatin structure.

**SUMMARY STATEMENT:** Nuclear blebs are linked to various pathologies, including cancer and premature aging disorders. We investigate alterations in higher-order chromatin structure within blebs, revealing fragmentation of nanoscopic heterochromatin domains.

## INTRODUCTION

The mammalian cell nucleus is a membrane-enclosed organelle that provides an enclosure for chromatin, the assembly of DNA and associated proteins that regulates critical processes such as gene transcription, replication, and DNA repair. Chromatin, chromatin proteins, and chromatin-related processes directly influence nuclear mechanics and shape.[1–5] Nuclear stability is further maintained by multiple processes, including by the nuclear lamina, a meshwork of type V intermediate filament proteins called lamins.[6] Besides its role in maintaining nuclear stiffness and stability, the lamina plays critical roles in regulating gene expression and DNA replication through chromatin interactions.[2] Located immediately underneath the inner nuclear membrane, the lamina consists of four major types of lamin proteins: lamin A, lamin C, lamin B1, and lamin B2. A-type lamins, which consist of lamins A and C, are primarily associated with developmental roles, contribute to nuclear stiffness, mainly expressed in differentiated cells, and are spatially located near the nucleoplasm.[1, 7] B-type lamins, in contrast, are expressed in all cell types throughout development and differentiation, provide global integrity of chromatin structure through chromatin-tethering, and are tightly associated with the inner nuclear membrane.[7–9] In mammalian cells, lamins interact with heterochromatin to form lamina-associated domains (LADs), identified through the DamID technique which maps protein-DNA interactions in a genome-wide manner, and are typically transcriptionally repressive environments.[10–12] Disruption of these LADs has been linked to epigenetic changes in cancer and pre-malignant processes such as the onset and evasion of senescence.[13]

Abnormal nuclear morphology and disruption of genome organization are associated with pathologies such as laminopathies (e.g., Hutchinson-Gilford progeria syndrome), cancer, and cardiac disorders.[2, 14, 15] Among the most radical transformations in nuclear shape is the protrusion of chromatin from the nuclear surface, known as a nuclear bleb, that are associated with pathological transformation.[2, 6, 15, 16] While these blebs are highly associated with gene-rich euchromatin and are believed to only contain lamin A/C,[7] recent evidence has indicated that non-canonical blebs also contain B-type lamins.[2, 17, 18] A- and B-type lamins both contribute to nuclear mechanics and morphology, and depletion of either has been widely shown to induce both abnormal nuclear shape and a higher propensity for nuclear rupture, increased presence of micronuclei, and more nuclear blebbing events.[14, 16, 19] Nucleus micromanipulation force measurements reveal that the nucleus is softer upon inhibition of either histone deacetylation (HDAC) or histone methyltransferase (HMT) which leads to nuclear bleb formation independent of lamins.[2, 15] Thus, chromatin and lamins resist external antagonistic forces from actin contraction[7, 20–22] and compression[23, 24], as well as internal transcription forces[4], to maintain nuclear shape. These studies indicate that nuclear mechanics are influenced by the balance of euchromatin and heterochromatin, and that perturbation of this balance can result in abnormal nuclear morphology and DNA damage, both hallmarks of human disease.[15, 16, 25] At the nuclear periphery, the dynamics of cytoskeleton reorganization and chromatin structural changes contribute to mechano-transduction and transcription, independent of lamins. For example, mechanosensitive ion channels embedded in the plasma membrane activate Ca^2+^ signaling upon cellular stress, which can contribute to heterochromatin reorganization and chromatin mobility.[15, 26–28]

Recent work demonstrates that chromatin assembles into higher-order polymeric domain structures (nanodomains, packing domains, cores), which range between 50-200 nm in size and contain ∼200 Kbp to 2Mbp of genomic content across multiple cell types.[29–31] A crucial feature of these domains is the formation of high-density centers with surrounding regions of decreased density until a transition into low-density space with RNA-polymerase activity forming primarily at the boundary. In the context of these findings, the structure of the genome assembles from disordered nucleosomes (5 to 25nm) transitioning into domains (50-150nm) and then into territorial polymers (>200nm). As has been previously shown, within the regime of chromatin assembling into domains, chromatin acts as a power-law polymer with dimension, *D* relating how the mass is distributed within the occupied volume. Notably, within supra-nucleosomal length scales, chromatin is not assembled purely as a space-fulling globule (D=3) nor is it as a poorly structured polymer with monomers primarily favors solvent interactions (D=5/3), instead it is typically within these ranges and varies from cell to cell. A key feature identified in this higher-order assembly is the coupling between heterochromatin centers (dense cores) with euchromatic periphery and a corrugated periphery [30]. As power-law polymeric assemblies, the structure of these chromatin packing domains is quantifiable by the relationship *M* ∝ *r^D^*, relating how genome content fills an occupied volume as a function of its radial distance, *r.* Notably, this organization can be directly measured by live-cell dual-Partial Wave Spectroscopic (dual-PWS) Microscopy and quantified by the fractal dimension, *D* **(Materials and Methods)**. In addition to quantifying packing domain structure in live cells, PWS microscopy allows measurement of the effective diffusion coefficient, *D_e_*, and the fractional moving mass (FMM) which quantifies the fraction of chromatin demonstrating coherent motion within a diffraction limited volume. Utilizing this technique, we have previously demonstrated that B-type lamin depletion is associated with increased levels of chromatin fractional moving mass and repositioning of heterochromatin cores [32, 33].

A major challenge in studying alteration in chromatin due to blebbing is that these represent infrequent, but critical events in nuclear structure. As such, they are difficult to assess using sequencing-based methods that measure ensemble chromatin organization such as Hi-C or ChIP-Seq and require the utilization of microscopic methods that can directly quantify changes in high-order genome structure. Therefore, in this study, we utilize live-cell PWS microscopy to investigate the interplay between the disruption of the nuclear lamina and heterochromatin enzymes in the structure of higher-order chromatin within blebs. Our results indicate distinct roles for the nuclear lamina and heterochromatin remodeling processes in regulating higher-order chromatin domains both of which are associated with bleb formation. Finally, pairing our findings with super-resolution microscopy, we show that a key transformation of higher-order chromatin within blebs is that of nanoscopic cores.

## RESULTS

### B-type lamin depletion or heterochromatin loss promote aberrant nuclear morphology in HCT-116 cells

Bleb formation has been identified in numerous cell types, but the frequencies of bleb formation have been shown to depend on multiple factors. Therefore, we first investigated the role of processes well established to induce bleb formation: inhibition of B-type lamins and disruption in heterochromatin enzymes.[2, 34] To assess the impact of lamin degradation on nuclear morphology, we applied the AID system to HCT116 colorectal carcinoma epithelial cells to induce simultaneous degradation of lamin B1 and lamin B2 as previously described.[32, 33, 35] Using immunofluorescence imaging, we quantified the percentages of nuclear blebbing in HCT116^LMN(B1&B2)-AID^ cells before and after depletion of B-type lamins by auxin treatment for 24 hours. We found that in comparison to untreated controls, auxin treatment promoted a significant increase in the percentage of cells containing nuclear blebs (2.07% vs 6.23%; 4.163 ± 1.033 (Mean difference ± SEM), p-value = 0.016, Student’s t-test) (**Fig. 1A; Fig. S1A**), in agreement with past studies. [2, 36, 37] Previous work from this group has demonstrated that inhibition of HDACs to increase euchromatin content in mammalian cells or inhibition of histone methyltransferases to decrease heterochromatin content results in a softer nucleus and promotes nuclear blebbing, without perturbing lamins.[2, 15] We therefore hypothesized that in addition B-type lamin loss increasing nuclear blebbing, heterochromatin loss would also result in a substantial increase in nuclear deformations in HCT-116 cells. To test this, we treated HCT116^LMN(B1&B2)-AID^ cells with either GSK343, an inhibitor of the histone methyltransferase Enhancer of Zeste Homolog 2 (EZH2), or Trichostatin A (TSA), an inhibitor of class I and II HDACs for 24 hours.

**Figure 1.**
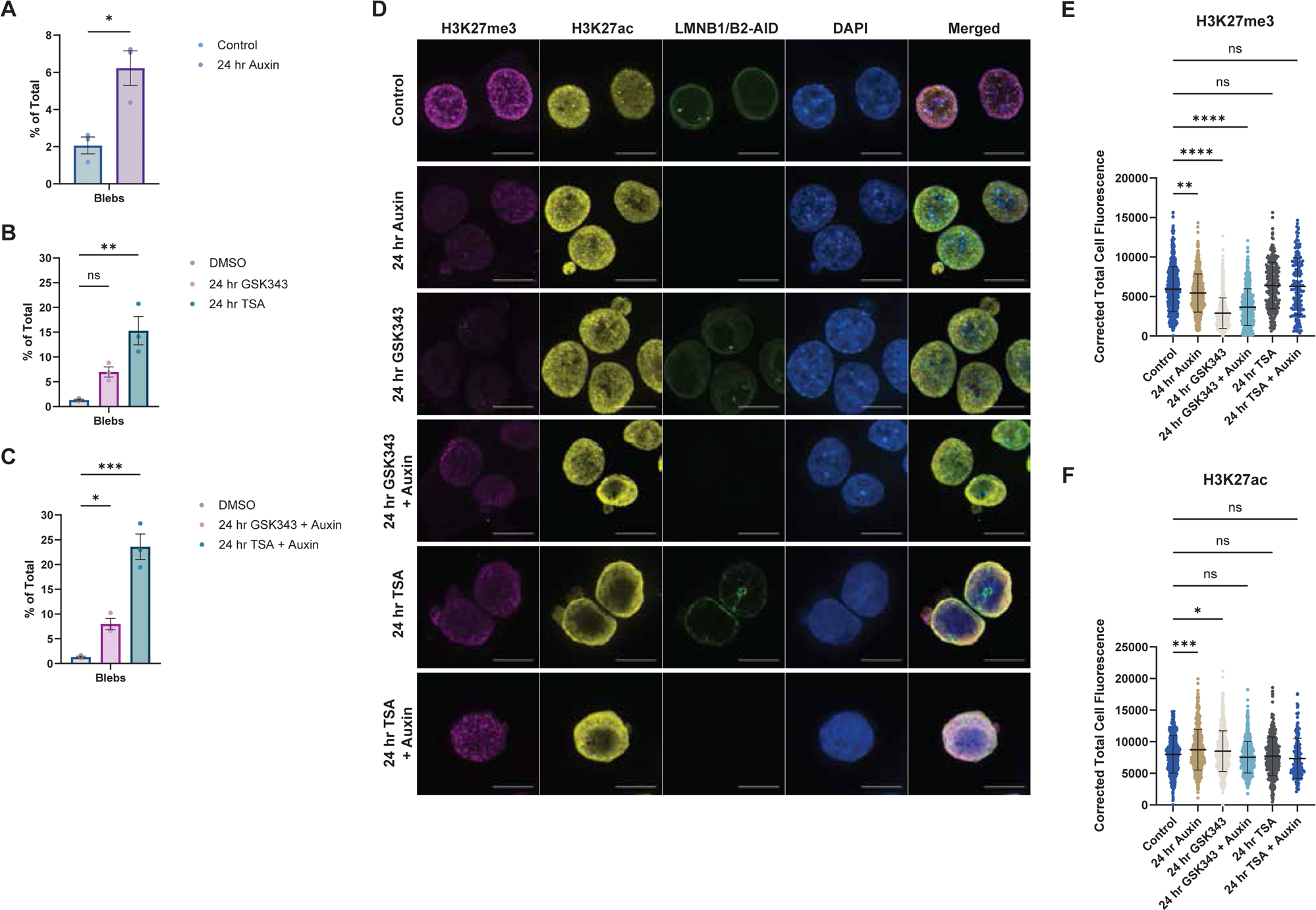
Aberrant nuclear morphology is induced by the loss of B-type lamins or heterochromatin. **(A)** Percentages of nuclear deformations compiled over each field of view for untreated (control) and 24-hour auxin conditions in HCT116^LMN(B1&B2)-AID^ cells. Each dot represents a technical replicate (N = 3; Control n = 1102, Auxin n = 1081). **(B)** Percentages of nuclear deformations compiled over each field of view for DMSO (vehicle control), 24-hour GSK343, and 24-hour TSA treatment conditions. Each dot represents a technical replicate (N = 3; DMSO n = 295, GSK343 n = 1096, TSA n = 456). **(C)** Percentages of nuclear deformations compiled over each field of view for DMSO (vehicle control), 24-hour GSK343 with auxin, and 24-hour TSA with auxin treatment conditions in HCT116^LMN(B1&B2)-AID^ cells. Each dot represents a technical replicate (N = 3; DMSO n = 295, GSK343 + Auxin n = 933, TSA + Auxin n = 456). For **(A-C)**, bar plots are represented as mean ± SEM. Unpaired two-tailed t-test with Holm-Šídák test for multiple comparisons applied in **(B-C)**. **(D)** Representative images of H3K27me3 (magenta), H3K27ac (yellow), Lamin B1/B2-AID (green), DAPI (blue), and merged fluorescence for control, 24-hour auxin, 24-hour GSK343, 24-hour GSK343 + Auxin, 24-hour TSA, and 24-hour TSA + Auxin treatment conditions in HCT116^LMN(B1&B2)-AID^ cells. Scale bar = 10 µm. **(E)** Corrected total cell fluorescence measurements of H3K27me3 and **(F)** H3K27ac for control, 24-hour auxin, 24-hour GSK343, 24-hour GSK343 + Auxin, 24-hour TSA, and 24-hour TSA + Auxin treatment conditions in HCT116^LMN(B1&B2)-AID^ cells. Each dot represents a cell nucleus. Violin plots show the median and quartiles. Error bars represent mean ± SD. One-way ANOVA with Dunnett’s test for multiple comparisons. For **(D-F)**, data are representative of two technical replicates (N = 2, total n > 150 for each condition). *P ≤ 0.05, **P ≤ 0.01, ***P ≤ 0.001, ****P ≤ 0.0001

Our results confirmed that GSK343 and TSA treatment significantly increased the percentage of blebbed nuclei within HCT-116 cells; with TSA treatment also inducing micronuclei formation and associated with severely deformed nuclear periphery (**Fig. 1B; Fig. S1A**). The effect of TSA treatment on nuclear blebbing frequency was drastically higher than that of auxin or GSK343 in HCT116^LMN(B1&B2)-AID^ cells (Auxin 6.23%, GSK343 6.99%, TSA 15.32%). Next, we explored how combined treatment of either auxin and GSK343 or auxin and TSA would impact the frequency of nuclear deformations. In contrast to prior work, combination of inhibition of b-type lamins with disruption of heterochromatin resulted in only slightly increased rates of nuclear deformations in comparison to lamin depletion, GSK343, or TSA treatment alone (Auxin + GSK343 7.97%, Auxin + TSA 23.56%) (**Fig. 1C; Fig. S1A**). Using immunofluorescence microscopy, we visualized these blebs and compared relative levels of H3K27me3 and H3K27ac between the untreated control and auxin, GSK343, and TSA-treated conditions within the cell nucleus (**Fig. 1D**). Consistent with prior reports, we observe a reduction in H3K27me3 levels upon auxin-induced degradation of B-type lamins, GSK343, and TSA treatment, however, only a concomitant increase in H3K27ac levels was observed in the auxin-treatment group (**Fig. 1E, F**). Taken together, these results indicated that TSA-mediated heterochromatin disruption promotes nuclear deformations to similar levels or greater levels than B-type lamin loss in HCT-116 cells.

### Nanoscale chromatin packing domains are disrupted within nuclear blebs

We recently demonstrated that the decreased DNA density is conserved across multiple bleb mechanisms and is a consistently preserved feature of blebs. We investigate here in greater detail the influence in the change of higher-order chromatin organization upon bleb formation. We have previously shown that live-cell PWS microscopy, which does not resolve each individual domain but measures the local ensemble in individual nuclei (**Materials and Methods**), is sensitive to detecting the assembly into supra-nucleosome structures in individual cells with measurements comparable to those observed on electron microscopy by measuring the variations in the visible-light interference spectrum from within the nucleus. [29, 30, 38–40] Likewise, we have shown that by analyzing the temporal interference spectrum at a single-wavelength dual-PWS microscopy can measure the temporal evolution of chromatin density and the fractional moving mass (FMM), which measures the volume fraction of- and mass of-chromatin moving coherently with a sensitivity to mass density fluctuations as low as ∼5*10^−21^ grams, and the effective diffusion coefficient (D_e_) within the nucleus (ranging between ∼0.065μM^2^/s to 3.5*10^−5^ μM^2^/s). In the context that the mass of an individual nucleosome ∼10^−19^ grams, the typical values of FMM measured represent the movement of nucleosome clutches moving coherently (as an ensemble). With respect to the D_e_, the observed values are typically between the observed rate of diffusion for genomic loci (∼10^−4^ μM^2^/s) and the rate of mRNA through the nucleus (∼5*10^−2^ μM^2^/s).[30] Given these considerations, we utilized dual-live cell PWS microscopy to probe the higher-order structure of chromatin in blebs, nuclei with blebs, and stable nuclei.

Applying dual-PWS to the three well known processes that contribute to bleb formation, we investigated the structure of higher order chromatin and mobility in B-type lamin depletion, HDAC inhibition, and in EZH2 inhibition. As each perturbation have a distinct means to promote bleb formation, we first evaluated the structure of chromatin packing domains observed within blebs in all conditions (controls, EZH2i, HDACi, and lamin depletion) to see if any commonalities were present (**Table 1**; **Fig. 2A-F**). Overall, this indicated that independent of the mechanism of bleb formation, relative to the nuclear body higher-order chromatin organization within blebs was associated with a lower-likelihood of well-formed packed domains (low *D*) and fragmented clutches (decreased FMM) with increased mobility (*D_e_).* Comparing the observed behavior of chromatin domains across the nuclei in these conditions, we observed that treatment with EZH2 or TSA resulted in decreased *D* with an associated increase in FMM in comparison to untreated controls whereas *D* and FMM increased upon B-type lamin depletion, indicating that disruption of heterochromatin enzymatic processes result in fragmentation of chromatin domains and the loss of coherent chromatin motion. Next, we compared the behavior of chromatin domains within blebs across all conditions and unexpectedly observed that domains associated due to lamin B depletion had a comparably higher *D* and FMM compared to those that occur due to inhibition of heterochromatin remodeling enzymes (**Fig. 2A, B, D, E; Fig. S1B, C**). As such, these results suggest that the domains translocating into blebs are larger in size and more stable with the disruption of the lamina compared to the other conditions. As removal of B-type lamins led to a significant increase in nuclear blebbing, these findings point to a bleb-associated chromatin phenotype, in which nuclear blebs contain fewer packing domains, and B-type lamin degradation could reduce the barriers for nuclear bleb formation that results from the restructuring or redistribution of packing domains.

**Figure 2.**
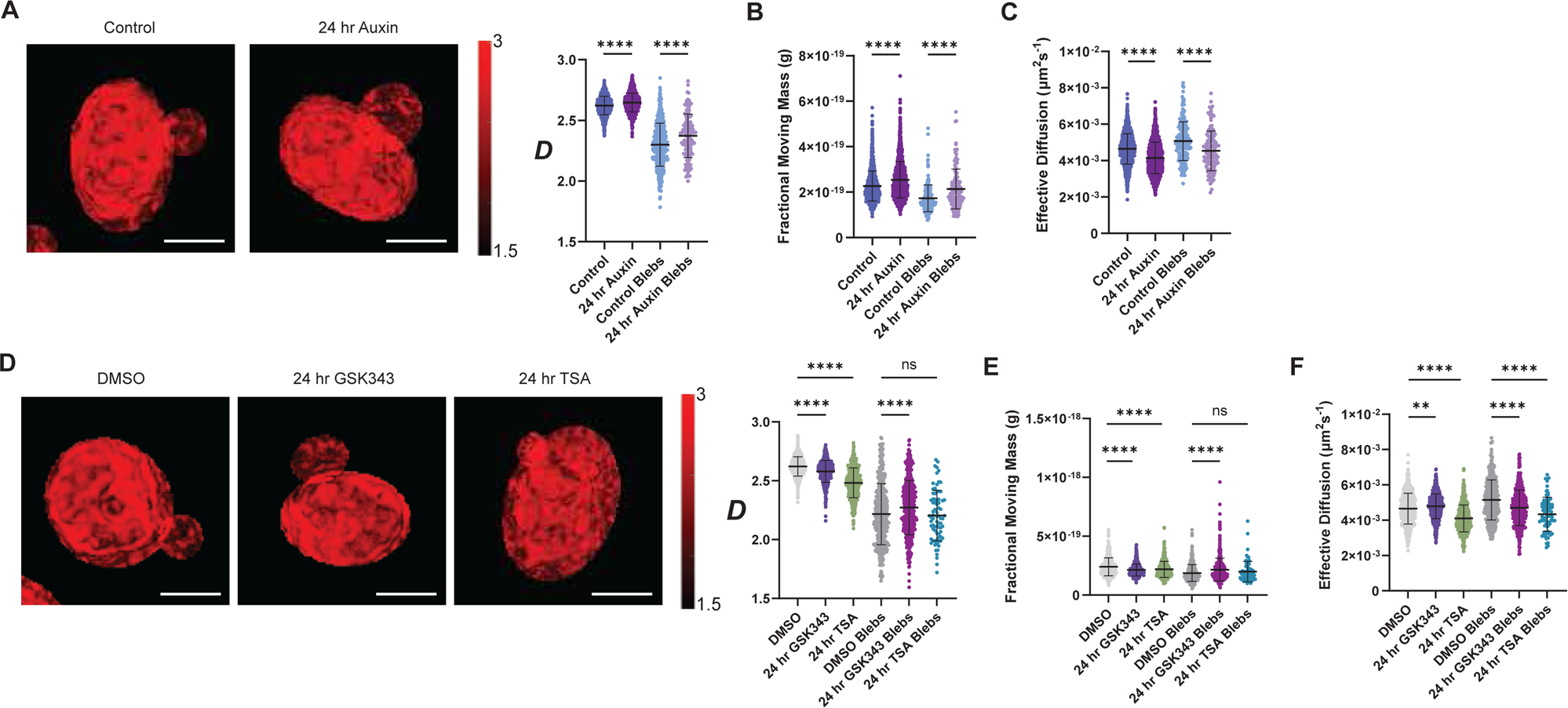
Loss of B-type lamins, EZH2i, and HDACi induce a bleb-associated chromatin phenotype. **(A)** Representative PWS *D* maps and *D* values for the nuclear bodies and nuclear blebs for control and 24-hour auxin-treatment conditions in HCT116^LMN(B1&B2)-AID^ cells. **(B)** Fractional moving mass values for nuclear bodies and nuclear blebs for control and 24-hour auxin-treatment conditions in HCT116^LMN(B1&B2)-AID^ cells. **(C)** Effective diffusion coefficient values for the nuclear bodies and nuclear blebs for control and 24-hour auxin-treatment conditions in HCT116^LMN(B1&B2)-AID^ cells. **(D)** Representative PWS *D* maps and *D* values for the nuclear bodies and nuclear blebs for DMSO (vehicle control), 24-hour GSK343, and 24-hour TSA-treatment conditions in HCT116^LMN(B1&B2)-AID^ cells. **(E)** Fractional moving mass values for the nuclear bodies and nuclear blebs for DMSO (vehicle control, 24-hour GSK343, and 24-hour TSA-treatment conditions in HCT116^LMN(B1&B2)-AID^ cells. **(F)** Effective diffusion coefficient values for the nuclear bodies and nuclear blebs for DMSO (vehicle control), 24-hour GSK343, and 24-hour TSA-treatment conditions in HCT116^LMN(B1&B2)-AID^ cells. For **(A-F)**, each dot represents a cell nucleus. (Control n = 2451, Auxin n = 2140, Control Blebs n = 200, Auxin Blebs n = 129, DMSO n = 741, GSK343 n = 790, TSA n = 498, DMSO Blebs n = 564, GSK343 Blebs n = 467, TSA Blebs n = 77). Error bars represent mean ± SD. Data are compiled from three technical replicates (N = 3). Violin plots show the median and quartiles for the unpaired two-tailed t-test between selected groups. *P ≤ 0.05, **P ≤ 0.01, ***P ≤ 0.001, ****P ≤ 0.0001. Scale bars = 5 µm.

**Table 1.** Dual-PWS measurements of nuclear blebs.

| Condition | Average *D*<br>(Mean ± SEM) | Average FMM (g)<br>(Mean ± SEM) | Average *D<sub>e</sub>* (μM<sup>2</sup>/s)<br>(Mean ± SEM) |
| --- | --- | --- | --- |
| Control (Untreated) | 2.30 | 1.76*10<sup>-19</sup> | 5.09*10<sup>-3</sup> |
| Control (DMSO) | 2.22 | 1.89*10<sup>-19</sup> | 5.12*10<sup>-3</sup> |
| **(-) Lamin B1/B2** | 2.37 (± 0.03) | 2.26*10<sup>-19</sup> (± | 4.35*10<sup>-3</sup> (± 0.0003) |
| **(-) EZH2** | 2.27 (± 0.03) | 2.16*10<sup>-19</sup> (± 1.53*10<sup>-20</sup>) | 4.71*10<sup>-3</sup> (± 0.0001) |
| **(-) HDACs** | 2.20 (± 0.03) | 1.99*10<sup>-19</sup> (± 1.53*10<sup>-20</sup>) | 4.34*10<sup>-3</sup> (± 0.0001) |

To further characterize the temporal dynamics of nuclear blebbing induced by either B-type lamin loss or heterochromatin disruption, we used dual-PWS to measure chromatin mobility between nuclear bleb and nuclear body in live cells. As TSA treatment resulted in the most substantial increase in the frequency of nuclear deformations above, we treated HCT116^LMN(B1&B2)-AID^ cells with either DMSO or TSA. By measuring the spectral interference at a single wavelength as a function of time from within nuclei and within blebs, we could directly image and measure how mass was transitioning between these phases.[39] As discussed previously, variations in the temporal interference quantifies the FMM while the spatially-average signal is inversely proportional the chromatin volume concentration (chromatin density).[39, 41] Utilizing this approach, we can visualize without labels the temporal evolution of chromatin density in both the nucleus and the bleb with a temporal resolution of 50ms per frame (acquired over 15s total in this instance). On imaging chromatin motion in the DMSO control nuclei, its visually apparent that density moves randomly over these timescales. In contrast, within a TSA treated cell with a visually apparent bleb, chromatin decreases rapidly adjacent to the nuclear bleb (**Fig. 3A, B; Movie S1-3)**. Between the nuclear bleb and the main nuclear body in TSA treatment, we observed the active transit of small amounts of mass as evidenced by increased local chromatin density and the visual changing of density in the bleb. Comparing the density of chromatin within DMSO associated and TSA induced blebs, TSA blebs unexpectedly had higher chromatin concentration compared to the DMSO control group (**Fig. 3C**). Our results therefore further suggest that the mobilization of chromatin packing domains is an active process during nuclear blebbing induced by B-type lamin and heterochromatin loss.

**Figure 3.**
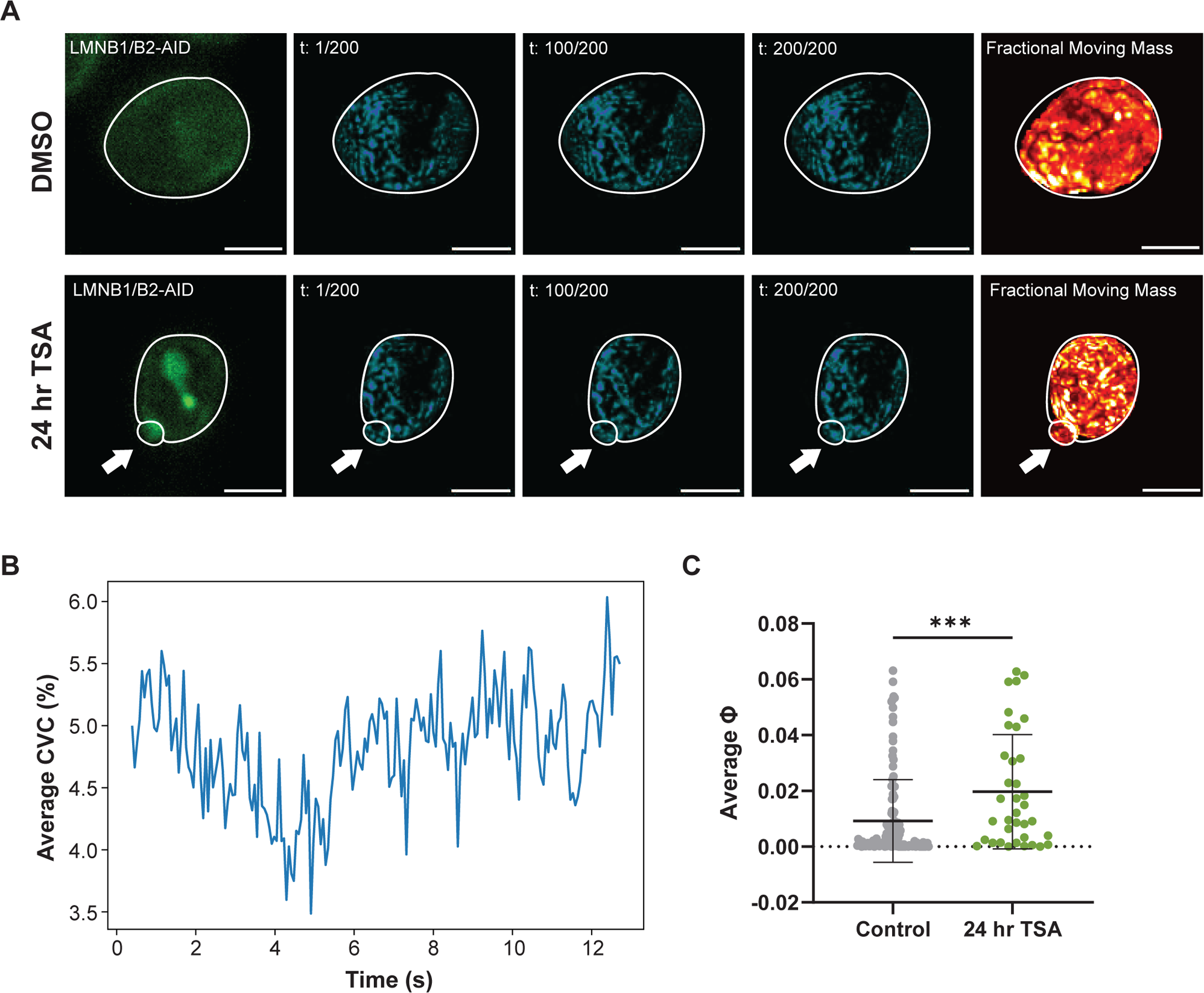
Chromatin density decreases rapidly in the boundary adjacent to the nuclear bleb. **(A)** Representative mClover signal, fractional moving mass “hot” heat map, and individual frames of the temporal interference signal, inversely proportional to chromatin density over the imaging acquisition time of live HCT116^LMN(B1&B2)-AID^ cells for DMSO (control) and 24-hour TSA treatment from Dual-PWS. Arrows indicate nuclear blebs. Data are representative of three technical replicates (N = 3; DMSO n = 741, TSA n = 498). Fluorescent, fractional moving mass, and individual frame images have the same scaling between treatment conditions. Scale bars = 5 µm. **(B)** Representative chromatin volume concentration (CVC) of a bleb over the acquisition time of dynamic PWS signal. The zig-zag behavior of the temporal signal originated from the moving nature of chromatin and is used to measure FMM. The general trend indicates mass moving out and then moving into the bleb. **(C)** Bleb average CVC for DMSO (control) and 24-hour TSA treatment measured from dynamics PWS signal. Data are the same as of (A). ***P ≤ 0.001

### Super-resolution imaging of chromatin heterochromatin domains in nuclear blebs

Nuclear lamins and heterochromatin have been shown to act in parallel to maintain the mechanical properties of the nucleus but the consequence of these on chromatin nanodomains in bleb formation have not been investigated.[28, 34, 42] Additionally, chromatin structure and dynamics are often closely related, which may support mechanisms of either granting or limiting access to regions with high local chromatin concentration[43]. In the context of prior work suggesting that chromatin domains are composed of high-density, presumably heterochromatic centers[29–31, 44], we investigated the transformation in constitutive heterochromatin domains between the nuclear bleb and the nuclear body in spontaneously forming blebs (controls), in lamin B1/B2 depletion-associated blebs, and in heterochromatin enzyme-inhibited blebs (TSA) using super-resolution microscopy. Due to the limitation of bleb formation being a low-frequency process (< 15% of the time), we were only able to identify blebs in a few nuclei in total in HCT-116 cells (**Fig. 4A-C**). Given this limitation, we utilized a second cell-line model, U2OS cells, that were associated with higher rates of bleb formation upon HDAC inhibition with TSA (**Fig. 4D-E**).

**Figure 4.**
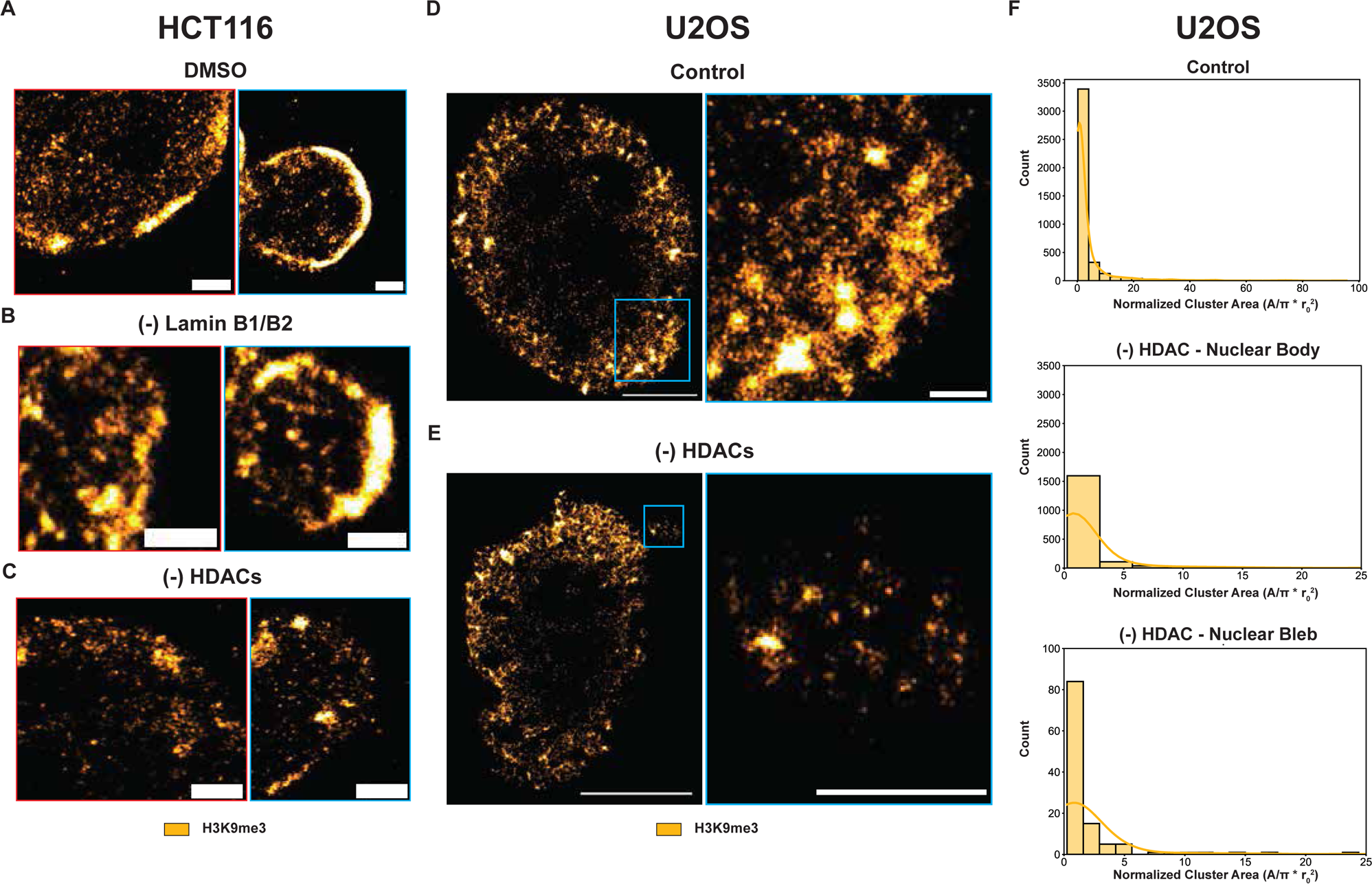
Heterochromatic nanodomains are reorganized during nuclear bleb formation. **(A)** Representative SMLM images of HCT116^LMN(B1&B2)-AID^ cells with zoomed-in views before and after 24-hour auxin treatment or 24-hour TSA treatment. Yellow: H3K9me3. Data are representative of three technical replicates (N = 3; Control n = 1, Auxin (- Lamin B1/B2) n = 1, TSA (- HDAC) n = 1). Scale bars = 5 µm for whole nucleus, 1 µm for inset of whole nucleus (red) and nuclear bleb (blue). **(B)** Representative SMLM images of U2OS cells with zoomed-in views before and 24-hour TSA treatment. Yellow: H3K9me3. Data are representative of three technical replicates (N = 3; Control n = 3, TSA (- HDAC) n = 3). Scale bars = 5 µm for whole nucleus, 1 µm for inset of nuclear bleb (blue). **(C)** Quantification of the number and size of heterochromatin nanodomains in control and TSA treatment conditions for U2OS cells.

Visually, we observed distinct differences in H3K9me3 chromatin nanodomains in these three conditions. In blebs formed spontaneously (**Fig. 4A**), blebs formed in B-type lamin depletion (**Fig. 4B**), and blebs formed due to inhibition of histone deacetylases (**Fig. 4C, E**), it is visually apparent that nanoscopic heterochromatin domains are observed. Within the nuclear body, as previously demonstrated[32, 33], auxin-induced depletion of B-type lamins resulted in reduced peripheral heterochromatic cores at the nuclear periphery, however, domains formed within the nuclear interior were typically larger in size (**Fig. S2**). With respect to heterochromatin nanodomains in GSK343 treated HCT116 cells[32, 33] and TSA treated U2OS cells, these were smaller than those in control cells and in lamin B-depletion as expected due to the inhibition of heterochromatin enzymes within the nuclear body (**Fig. 4D-F; Fig. S2**). In contrast to prior work, we found that the nuclear blebs arising from either B-type lamin degradation or TSA treatment contained heterochromatin around the periphery of blebs and within the center of the bleb (**Fig. 4C, E, F; Fig. S2**). This finding highly contradicts the plethora of research stating that all nuclear blebs are devoid of heterochromatin.[45–47] Instead, these results show that independent B-type lamin loss, HDAC inhibition gives rise to non-canonical nuclear blebs enriched in heterochromatin around near their boundaries and the transfer of nanoscopic heterochromatin domains into the bleb. This also challenges the notion that euchromatin enrichment is the most reliable marker of nuclear blebs[17], and further suggests that other cellular mechanisms could play a role in the morphological properties of these herniations.

## DISCUSSION

In this work, we found that nuclear packing domains are transformed within blebs induced by the loss of either B-type lamins or inhibition of heterochromatin enzymes. Specifically, the domains observed within blebs were typically poorly formed, with increased fragmentation and a higher effective diffusion coefficient compared to the domains observed in the nuclear body independent of the conditions. Despite the conserved differences across groups, we notably saw that in domains associated with lamin B depletion, these were larger than those produced by inhibition of heterochromatin remodeling enzymes (GSK343 inhibition of EZH2 and TSA inhibition of HDACs) suggesting that the barrier to movement of domains or nucleosome clutches is larger in the loss of b-type lamins whereas the inhibition of heterochromatin enzymes fragments domains to facilitate deformations in the nuclear border (**Fig. 2A, D**). Given these findings, one possible and interesting explanation is that domains or clutches of nucleosomes move in concert into blebs through transiently evolving defects in the nuclear lamina. As such, depletion of b-type lamins may increase the frequency of barrier disruption events or potentially result in larger transient defects that allow passage of larger domains into the bleb body. Further supporting these findings were the observation that the structure of heterochromatin domains upon B-type lamin depletion are larger in size compared to those in nuclei treated with heterochromatin enzyme inhibitors on super-resolution microscopy (**Fig. 4A, B).** Likewise, although limited the low frequency of bleb events, domains observed within all blebs were smaller in size and more disperse than those observed in the adjacent nuclear body (**Fig. 4A, B**).

The changes observed in chromatin domains within blebs could be related to functional consequences in signaling, possibly arising from applied mechanical stress when the nuclear lamina or heterochromatin are disrupted. For example, using single-nucleus isolation and micromanipulation assays, we previously demonstrated that nuclei with reduced heterochromatin levels are softer and succumb to nuclear blebbing, while nuclei with more heterochromatin levels are stiffer and resist blebbing.[2] This chromatin histone-modification-based nuclear rigidity could be related to the differential transcriptional responsiveness (i.e., transcriptional plasticity) previously observed in low-versus high-chromatin packing areas upon exposure to external stressors.[48] In many cases, nuclear blebbing is a marker of cell death (i.e., apoptosis) and is often observed during normal developmental processes or in response to various extracellular stressors. The transformation of domains within blebs upon either B-type lamin or heterochromatin enzyme disruption could potentially accelerate these processes by increasing DNA damage or cytoskeletal reorganization. Alternatively, bleb formation could be a necessary event to maintain the stability of the remaining chromatin domains to ensure their continued optimal function by maintaining the optimal conditions for remaining domains to function.

Although loss of B-type lamins and inhibition of heterochromatin enzymes both induced nuclear morphological changes and increased FMM within nuclear blebs, it is important to consider that these perturbations may not always reflect the same changes in cell phenotype. While B-type lamins are required for proper spatial positioning of heterochromatin and gene-specific loci[8, 32, 33], B-type lamin loss and heterochromatin disruption may impact different cellular mechanisms that give rise to these morphological changes. For example, in previous work, we found that decreasing heterochromatin promoted decreased nuclear rigidity and increased nuclear blebbing without necessarily altering lamins.[2] Conversely, removal of B-type lamins resulted in both a reduction of heterochromatin and increased nuclear blebbing.[8, 32, 33] Therefore, while lamins and heterochromatin interact, depletion of either could have differential effects on chromatin organization. In this work, we also found that HDAC or EZH2 inhibition promoted more nuclear deformations in comparison to auxin treatment to remove B-type lamins. As combined treatment of either auxin and TSA or auxin and GSK343 did not result in a more significant increase in these deformations in comparison to TSA or GSK343 alone, our results support previous findings that conclude disruption of chromatin alone is sufficient to cause nuclear blebbing[2]. However, the distortions in the nucleus caused in part by the breakdown of connections between chromatin and the nuclear lamina may be intensified by pressure gradients resulting from external influences.[49]

These external factors, such as confinement imposed by the actin cytoskeleton or the surrounding environment, could further contribute to the increased deformation of the nucleus. Additionally, processes such as HDACi or HMTi could act by expanding the volume of heterochromatin centers or destabilizing packing domains altogether. In theory, as weak, unstable packing domain (i.e., nascent domain) cores expand in size, one possible consequence could be increased variations in temporally active processes, such as gene transcription, resulting in amplified chromatin motion. Consequently, modifications to higher-order chromatin assemblies could promote bleb formation by degrading packing domains and/ or altering chromatin-based nuclear mechanics. However, further assessment is needed to confirm this theoretical interplay between packing domain formation, nuclear mechanics, and transcription.

The complexity of interactions within the genome results in varying chromatin dynamics at different length scales. Intrinsic characteristics of chromatin, which involve the dynamic rearrangement of histones, interactions among chromosome segments, chromatin remodelers, replication proteins, and transcriptional regulators are required for proper spatiotemporal genome organization. Other than Dual-PWS, several techniques have been utilized to investigate the contributions of chromatin dynamics to this organization. For example, a combination of photoactivated localization microscopy (PALM) and tracking of single nucleosomes was recently applied to assess nucleosome-nucleosome interactions and cohesin-RAD21 in domain formation and dynamics.[50] In line with our results, TSA treatment increased chromatin dynamics. Recently, proximity ligation-based chromatin assembly assays have been applied to investigate the kinetics of nuclear lamina binding to newly replicated DNA in mouse embryonic fibroblasts.[51] Finally, computational models have been applied to probe the time evolution of the chromatin over the G1 phase of the interphase in *Drosophilla* that successfully predict dynamic positioning of all LADs at the nuclear envelope.[52] While Chromatin Scanning Transmission Electron Microscopy (ChromSTEM) does not have live-cell imaging capabilities to resolve chromatin mobility[29], future work may involve this high-resolution imaging technique to investigate how the shift in chromatin dynamics seen here could be related to shifts in chromatin density, volume, and shape.

## CONCLUSION

The formation of nuclear blebs is believed to be associated with aberrant gene expression in pathological conditions; nevertheless, our understanding of chromatin structure within blebs and the mechanisms resulting in their formation remain poorly understood. The biophysical characteristics of cell nuclei, including their mechanical properties and architecture, play a crucial role in shaping cell phenotype, shape, and function. Our research demonstrates that the transformation of chromatin nanoscopic packing domains may contribute to bleb formation given the structures seen within nuclear blebs on live-cell nanoscopic imaging and super resolution microscopy. This indicates that histone modifications converge in altering chromatin packing domains with the resulting change in structure influencing nuclear mechanics and morphology. As blebs are associated with increased DNA damage, it highlights the need for further investigation into how the change in chromatin structure (both nucleosome modifications and higher-order domains) contribute to transformation of chromatin function within bleb compartments. Future work further investigating blebs can help us understand what happens to gene-transcription within these deformations and how the reintegration of these components of the genome happen during mitosis. Finally, future work to decouple how additional bleb-promoting mechanisms (compression, contraction, translocation) simultaneously at different length scales to organize the genome remains open for further investigation.

## MATERIALS AND METHODS

### HCT116 Cell Culture

HCT116^LMN(B1&B2)-AID^ cells and U2OS cells were grown in McCoy’s 5A Modified Medium (#16600-082, Thermo Fisher Scientific, Waltham, MA) supplemented with 10% FBS (#16000-044, Thermo Fisher Scientific, Waltham, MA) and penicillin-streptomycin (100 μg/ml; #15140-122, Thermo Fisher Scientific, Waltham, MA). To create these cells, HCT116 cells (ATCC, #CCL-247) were tagged with the AID system as previously described.[32, 33] All cells were cultured under recommended conditions at 37°C and 5% CO_2_. All cells in this study were maintained between passage 5 and 20. Cells were allowed at least 24 h to re-adhere and recover from trypsin-induced detachment. All imaging was performed when the surface confluence of the dish was between 40–70%. All cells were tested for mycoplasma contamination (ATCC, #30-1012K) before starting perturbation experiments, and they have given negative results.

### Auxin Treatment

HCT116^LMN(B1&B2)-AID^ cells were plated at 50,000 cells per well of a 6-well plate (Cellvis, P12-1.5H-N). To induce expression of OsTIR1, 2 μg/ml of doxycycline (Fisher Scientific, #10592-13-9) was added to cells 24 hours prior to auxin treatment. 1000 μM Indole-3-acetic acid sodium salt (IAA, Sigma Aldrich, #6505-45-9) was solubilized in RNase-free water (Fisher Scientific, #10-977-015) before each treatment as a fresh solution and added to HCT116^LMN(B1&B2)-AID^ cells.

### GSK343 Treatment

HCT116^LMN(B1&B2)-AID^ cells were plated at 50,000 cells per well of a 6-well plate (Cellvis, P12-1.5H-N). Cells were given at least 24 hours to re-adhere before treatment. GSK343 (Millipore Sigma, #SML0766) was dissolved in DMSO to create a 10 mM stock solution. This was further diluted in complete cell media to a final treatment concentration of 10 µM.

### Trichostatin A (TSA) Treatment

HCT116^LMN(B1&B2)-AID^ cells were plated at 50,000 cells per well of a 6-well plate (Cellvis, P12-1.5H-N). Cells were given at least 24 hours to re-adhere before treatment. TSA (Millipore Sigma, #T1952) was diluted in complete cell medium and added to cells at a final treatment concentration of 300 nM.

### Immunofluorescence Sample Preparation

HCT116^LMN(B1&B2)-AID^ cells at a low passage (<P10) were plated at 100,000 cells per well of a 6-well glass-bottom plate (Cellvis, #P06-1.5H-N). Following auxin treatment, cells were washed twice with 1x Phosphate Buffered Saline (PBS) (Gibco, #10010031). Cells were fixed with 4% paraformaldehyde (PFA) (Electron Microscopy Sciences, #15710) for 10 minutes at room temperature, followed by washing with PBS 3 times for 5 minutes each. Cells were permeabilized using 0.2% TritonX-100 (10%) (Sigma-Aldrich, #93443) in 1x PBS, followed by another wash with 1x PBS for 3 times for 5 minutes each. Cells were blocked using 3% BSA (Sigma-Aldrich, #A7906) in PBST (Tween-20 in 1x PBS) (Sigma-Aldrich, #P9416) at room temperature. The following primary antibodies were added overnight at 4°C: anti-H3K27ac (Abcam, #ab177178, dilution 1:7000) and anti-H3K27me3 (Abcam, #ab6002, dilution 1:200). Cells were washed with 1x PBS 3 times for 5 minutes each. The following secondary antibodies were added for 1 hour at room temperature: Goat anti-Rabbit IgG (H+L) Alexa Fluor 568 (Abcam, #ab175471, dilution 1:1000) and Goat anti-Mouse IgG (H+L) Highly Cross-Adsorbed Secondary Antibody, Alexa Fluor Plus 647 (Thermo Fisher Scientific, #A32728, dilution 1:200). Cells were washed with 1x PBS 3 times for 5 minutes each. Finally, cells were stained with DAPI (Thermo Fisher Scientific, #62248, diluted to 0.5 μg/mL in 1x PBS) for 10 minutes at room temperature. Prior to imaging, cells were washed with 1x PBS twice for 5 minutes each.

### Immunofluorescence Imaging

Live and fixed cells were imaged using the Nikon SoRa Spinning Disk confocal microscope equipped with a Hamamatsu ORCA-Fusion Digital CMOS camera. Live cells were imaged under physiological conditions (37°C and 5% CO_2_) using a stage top incubator (Tokai Hit). Images were collected using a 60x/ 1.42 NA oil-immersion objective mounted with a 2.8x magnifier. mClover was excited with a 488 nm laser, Alexa Fluor 647 was excited with a 640 nm laser, and DAPI was excited with a 405 nm laser. Imaging data were acquired by Nikon acquisition software.

### Dual-PWS Imaging

For live-cell measurements, cells were imaged and maintained under physiological conditions (5% CO^2^ and 37°C) using a stage-top incubator (In Vivo Scientific, Salem, SC; Stage Top Systems). Live-cell PWS measurements were obtained using a commercial inverted microscope (Leica, DMIRB) using a Hamamatsu Image-EM charge-coupled device (CCD) camera (C9100-13) coupled to a liquid crystal tunable filter (LCTF, CRi VariSpec) to acquire monochromatic, spectrally resolved images ranging from 500-700 nm at 2-nm intervals as previously described.[38–40] Broadband illumination was provided by a broad-spectrum white light LED source (Xcite-120 LED, Excelitas). The system is equipped with a long pass filter (Semrock BLP01-405R-25) and a 63x oil immersion objective (Leica HCX PL APO). All cells were given at least 24 hours to re-adhere before treatment (for treated cells) and imaging. Briefly, PWS measures the spectral interference signal resulting from internal light scattering originating from nuclear chromatin. This is related to variations in the refractive index (RI) distribution and is captured by the microscope by calculating the standard deviation of the spectral interference at each pixel (Σ). Chromatin packing scaling *D* was then calculated using maps of Σ. Although it is a diffraction-limited imaging technique, PWS can measure chromatin packing behaviors because the RI of chromatin is proportional to the local density of macromolecules (e.g., DNA, RNA, proteins). PWS senses the complex inhomogeneous RI distribution of chromatins with length scale sensitivity around 20 – 200 nm, and associated it with fractal coefficient, as previously described.[30, 39, 41, 48] PWS measurements were normalized by the reflectance of the glass medium interface (i.e., to an independent reference measurement acquired in a region lacking cells on the dish). This allows acquisition of the interference signal that is directly related to RI fluctuations within the cell. Changes in *D* resulting from each condition are quantified by averaging cells, taken across 3 technical replicates. Average *D* was calculated by first averaging *D* values from PWS measurements within each cell nucleus and then averaging these measurements over the entire cell population for each treatment condition.

### Dynamic PWS Measurements

Temporal PWS data was acquired as previously described.[32, 39] Briefly, dynamics measurements (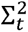, fractional moving mass (*m_f_*), and diffusion) are collected by acquiring multiple backscattered wide-field images at a single wavelength (550□nm) over time (acquisition time), to produce a three-dimensional image cube, where 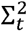 is temporal interference and t is time. Diffusion is extracted by calculating the decay rate of the autocorrelation of the temporal interference as previously described.[39] The fractional moving mass is calculated by normalizing the variance of 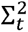 at each pixel. Using the equations and parameters supplied and explained in detail in the supplementary information of our recent publication [39], the fractional moving mass is obtained by using the following equation to normalize 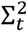 by *ρ*_0_ the density of a typical macromolecular cluster:

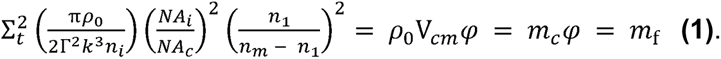

With this normalization, 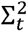 is equivalent to *m_f_*, which measures the mass moving within the sample. This value is calculated from the product of the mass of the typical moving cluster (*m_c_*) and the volume fraction of mobile mass (*φ*). *m_c_* is obtained by *m_c_* = V*_cm_ρ*_0_, where V*_cm_* is the volume of the typical moving macromolecular cluster. To calculate this normalization, we approximate *n_m_* = 1.43 as the refractive index (RI) of a nucleosome, *n*_1_ = 1.37 as the RI of a nucleus, *n_i_* = 1.518 as the refractive index of the immersion oil, and *ρ*_0_ = 0.55 g *cm*^−3^ as the dry density of a nucleosome. Additionally, *k* = 1.57E5 cm^−1^ is the scalar wavenumber of the illumination light, and Γ is a Fresnel intensity coefficient for normal incidence. *NA_c_* = 1.49 is the numerical aperture (NA) of collection and *NA_i_* = 0.52 is the NA of illumination. As stated previously[39], 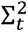 is sensitive to instrument parameters such as the depth of field, substrate refractive index, etc. These dependencies are removed through normalization with the proper pre-factor calculated above for obtaining biological measurements. It should also be noted that backscattered intensity is prone to errors along the transverse direction[39]. Due to these variations, these parameters are more accurate when calculating the expected value over each pixel.

Chromatin volume concentration is calculated by Fresnel reflection coefficient. Recall that the reflectance at a RI mismatch interface:

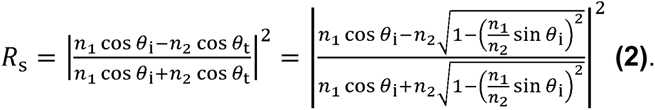

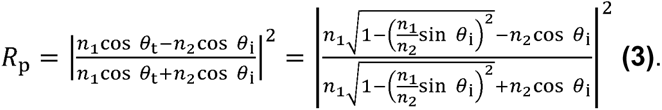

Although the chromatin is inhomogeneous and not infinitely large, the correlation between chromatin average RI and reflection coefficients still holds as confirmed by Finite Difference Time Domain (FDTD) simulations (**Fig. S3**). Briefly, we used home-built FDTD software to simulate the entire PWS imaging system, from incident to light-matter interaction and then collection.[53] Light beams representing the characteristics of experimental *NA_i_* are first introduced to the simulation space. The simulation space contains a layer of glass and random media that represents chromatin average RI and packing behavior. The EM wave after light-matter interaction is then collected with the same *NA_c_* and far field PWS image is analyzed the same way in experiments. We did a series of simulations with different average RI and measured the mean reflection coefficients. We have fitted:

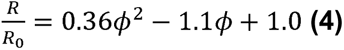

where *ϕ* is chromatin volume concentration, related to RI through Gladstone-Dale equation [48]:

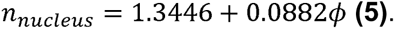

### SMLM Sample Preparation and Imaging

Primary antibody rabbit anti-H3K9me3 (Abcam, #ab176916, dilution 1:2000) was aliquoted and stored at –80°C. The secondary antibody goat anti-rabbit AF647 (Thermo Fisher Scientific, #A-21245, dilution 1:1000) was stored at 4°C. The cells were plated on No. 1 borosilicate bottom eight-well Lab-Tek Chambered cover glass with at a seeding density of 1.25 × 10^4^. After 48 hours, the cells were fixed in 3% paraformaldehyde in PBS for 10 minutes, and then subsequently washed with PBS once for five minutes. Thereafter the samples were quenched with freshly prepared 0.1% sodium borohydride in PBS for 7 minutes and rinsed with PBS three times at room temperature. The fixed samples were permeabilized with a blocking buffer (3% bovine serum albumin (BSA), 0.5% Triton X-100 in PBS) for 20 minutes and then incubated with rabbit anti-H3K9me3 in blocking buffer for 1-2 hours at room temperature and rinsed with a washing buffer (0.2% BSA, 0.1% Triton X-100 in PBS) three times. The fixed samples were further incubated with the corresponding goat secondary antibody–dye conjugates, anti-rabbit AF647, for 40 minutes, washed thoroughly with PBS three times at room temperature and stored at 4°C. Imaging of samples was performed on a STORM optical setup built on the commercially available Nikon Ti2 equipped with a Photometric 95B sCMOS camera and a 1.49 NA 100X oil immersion objective lens. Samples were illuminated with the MPB Communications 2RU-VFL-P-2000-647-B1R 647 nm 200 mW laser. Image acquisition was performed at 20-30 ms exposure for 10-15k frames.

### Data and Image Analysis

We used GraphPad Prism 10.1.1 for making all plots. For immunofluorescence imaging, maximum intensity projection of Z-series images was performed using FIJI.[54] To quantify nuclear deformation frequency, we considered blebs to be herniations that were still connected to the nuclear body. We considered ruptures to be cells that were no longer intact, and we considered micronuclei to be herniations that were no longer connected to the nuclear body and of similar sizes to nuclear blebs. For each field of view, the number of nuclei and the number of each nuclear deformation type was manually counted using FIJI. We then determined the percentages of total cells within each tested condition that displayed each nuclear deformation type.

### Super-resolution Data Analysis

We used the Thunder-STORM FIJI Plug-in [55]to apply Maximum Likelihood Estimation fitting of a gaussian point spread function to our image stack. Localization datasets were then put into our Python script that utilized DBSCAN (epsilon=50, min_pts=3) to cluster our localized heterochromatic events. Heterochromatin domain size was estimated by fitting a polygon to the peripheral cluster points using the *scipy* Convex Hull method. Outlier clusters smaller than twice the mean uncertainty of our localization (∼ 25 nm) or larger than 800 nm were removed from the analysis. Results displayed are concatenations of identified heterochromatic domains across all cells in that condition.

### Statistical Analysis and Quantification

Statistical analysis was performed using GraphPad Prism 10.1.1 and Microsoft Excel. Pairwise comparisons were calculated on datasets consisting of, at a minimum, biologically independent duplicate samples using a two-tailed unpaired t test or Mann-Whitney test. The type of statistical test is specified in each case. Experimental data are presented either the mean ± SEM or mean ± SD, as stated in figure legends. A *P* value of < 0.05 was considered significant. Statistical significance levels are denoted as follows: n.s. = not significant; \**P*<0.05; \*\**P*<0.01; \*\*\**P*<0.001; \*\*\*\**P*<0.0001. Sample numbers (# of nuclei, n), the number of replicates (N), and the type of statistical test used is indicated in figure legends.

## ACKNOWLEDGEMENTS

Microscopy was performed at the Biological Imaging Facility at Northwestern University (RRID:SCR_017767), graciously supported by the Chemistry for Life Processes Institute, the NU Office for Research, the Department of Molecular Biosciences and the Rice Foundation. FDTD simulations were performed at Quest, high-performance computing cluster at Northwestern University.

## DATA AVAILABILITY

All relevant data can be found within the article and its supplementary information. The cell lines have been authenticated and are available upon request. Further information and requests for resources and reagents should be directed to and will be fulfilled by the lead contact, Vadim Backman.

## AUTHORS’ CONTRIBUTIONS

E.M.P. and L.A. wrote the paper and performed immunofluorescence and Dual-PWS imaging and analysis. N.A. and L.A. conducted SMLM imaging and analysis. N.A. assisted with representative Dual-PWS images. Y.S. conducted density estimations from Dual-PWS data. R.G. set up the optical system for all super resolution image acquisition. E.M.P, A.S., L.A., and V.B. conceptualized the project and edited the manuscript.

## DIVERSITY AND INCLUSION STATMENT

One or more of the authors of this paper self-identifies as an underrepresented minority.

## SUPPLEMENTARY FIGURE TITLES AND LEGENDS

**Figure S1.**
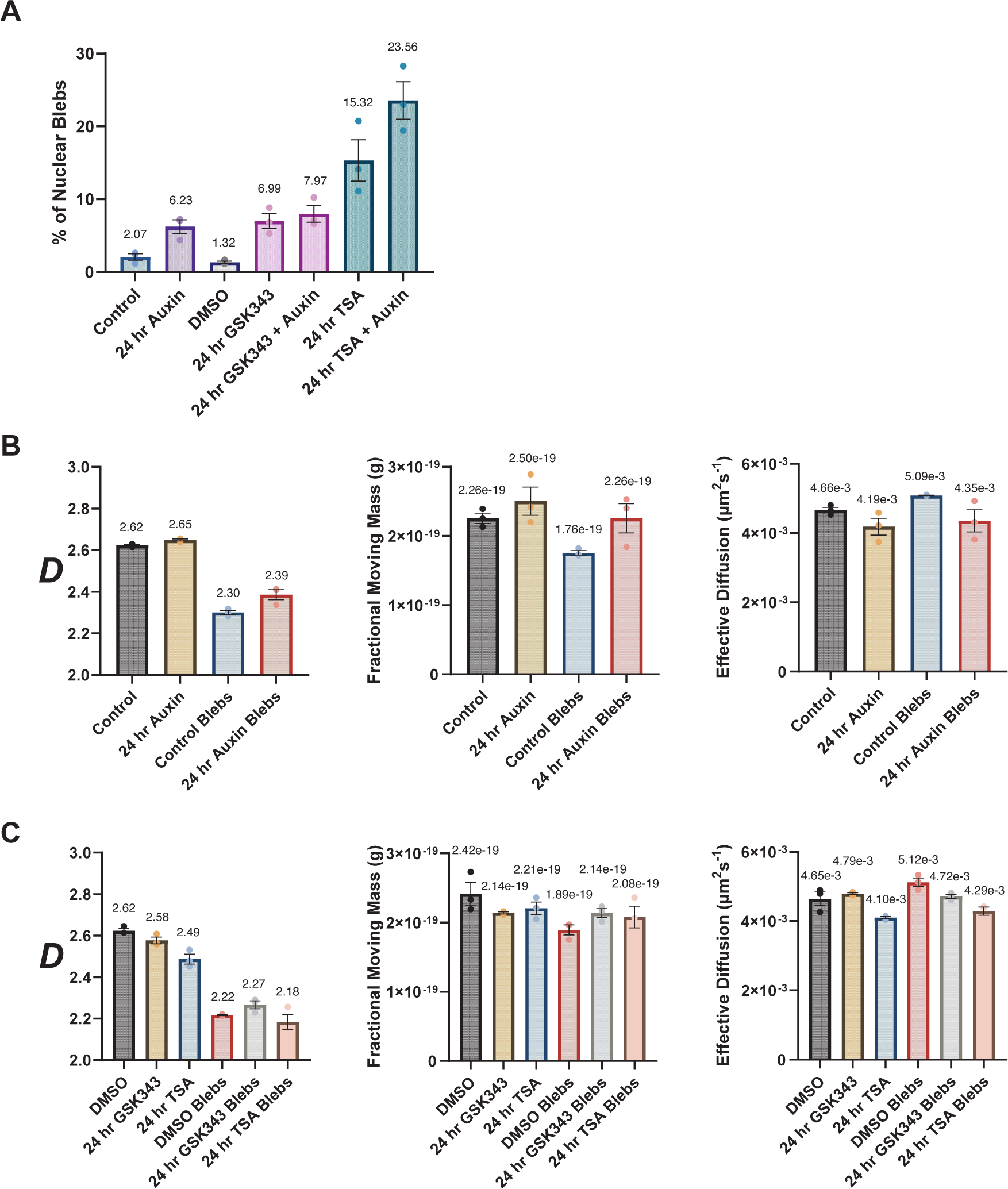
Nuclear blebbing induces the redistribution and reorganization of packing domains. **(A)** Averages of all three replicates for percentages of nuclear deformations compiled over each field of view for control, 24-hour auxin, DMSO (vehicle control), 24-hour GSK343, 24-hour TSA, 24-hour GSK343 with auxin, and 24-hour TSA with auxin treatment conditions. Each dot represents a technical replicate (N = 3; Control n = 1102, Auxin n = 1081, DMSO n = 295, GSK343 n = 1096, TSA n = 456, GSK343 + Auxin n = 933, TSA + Auxin n = 456). **(B)** Averages of all three replicates for *D* values, fractional moving mass, and effective diffusion coefficient for the nuclear bodies and nuclear blebs for control and 24-hour auxin-treatment conditions in HCT116^LMN(B1&B2)-AID^ cells. **(C)** Averages of all three replicates for *D* values, fractional moving mass, and effective diffusion coefficient for DMSO (vehicle control) 24-hour GSK343, and 24-hour TSA-treatment conditions in HCT116^LMN(B1&B2)-AID^ cells. For **(A-C)** means of all replicates are presented above each bar in the plots. Error bars represent mean ± SEM. For **(B-C)**, Data are compiled from three technical replicates (N = 3; Control n = 2451, Auxin n = 2140, Control Blebs n = 200, Auxin Blebs n = 129, DMSO n = 741, GSK343 n = 790, TSA n = 498, DMSO Blebs n = 564, GSK343 Blebs n = 467, TSA Blebs n = 77).

**Figure S2.**
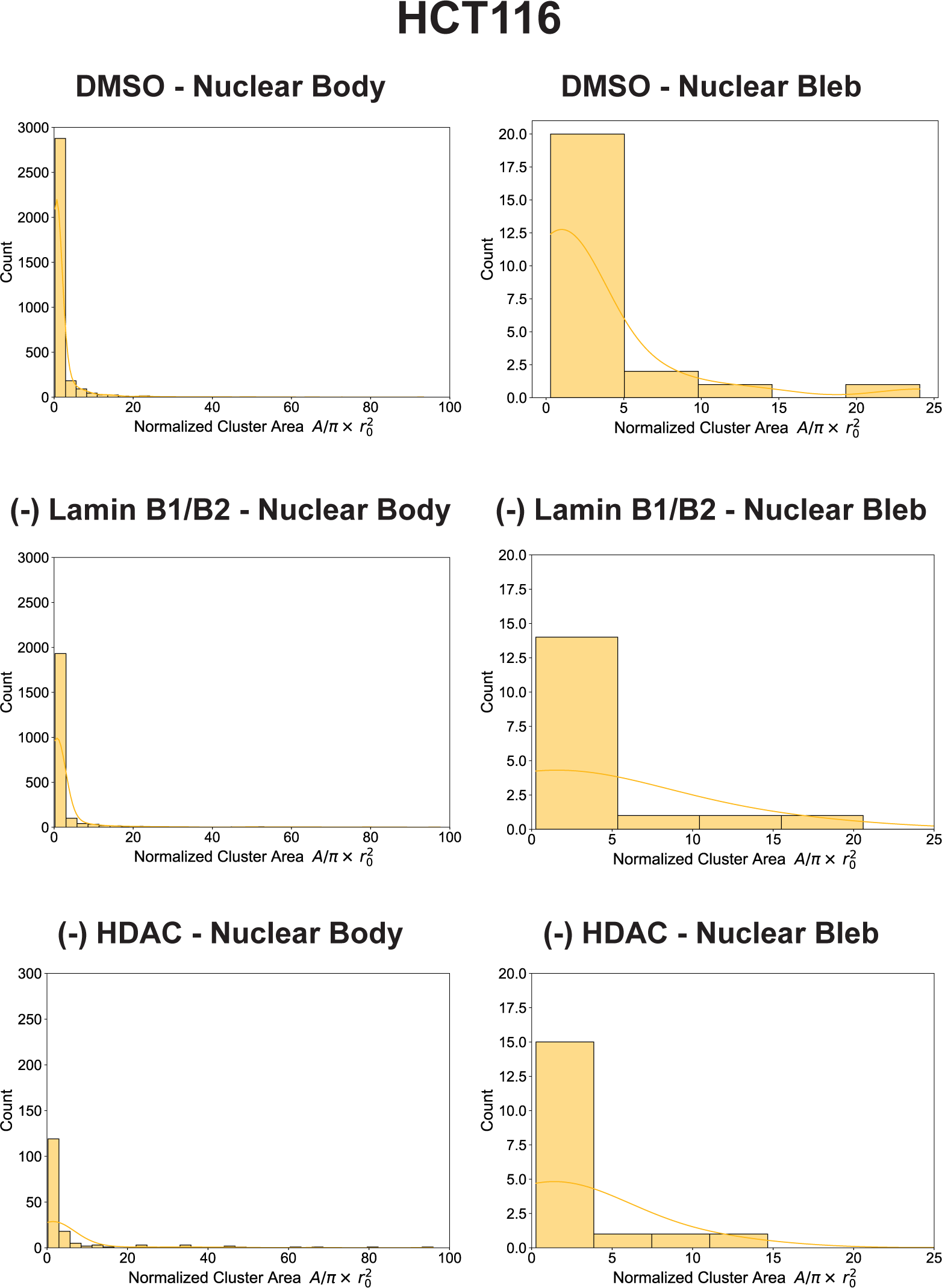
Characterization of heterochromatin nanodomains from SMLM images. Quantification of the number and size of heterochromatin nanodomains in control, loss of lamin B1/B2, and TSA treatment conditions for HCT116^LMN(B1&B2)-AID^ cells.

**Figure S3.**
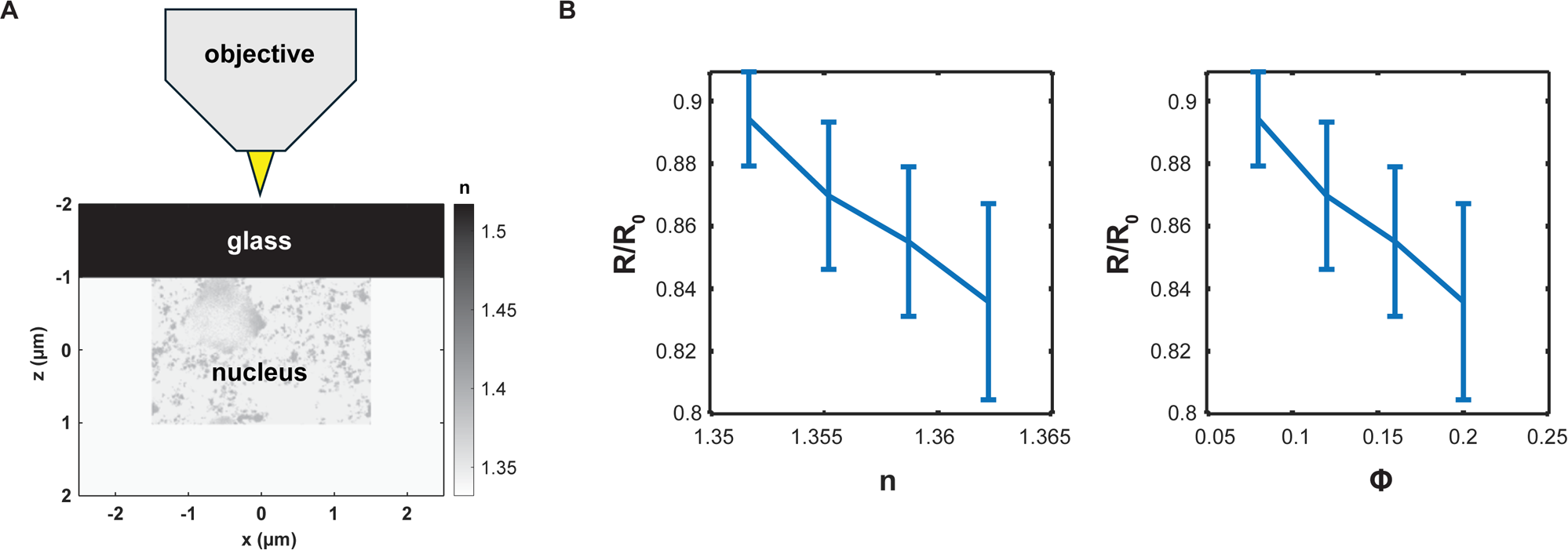
FDTD simulation confirms the correlation between PWS Intensity with nuclear average RI and CVC. **(A)** Schematic of FDTD simulation setup. Light is illuminated from objective and focused on the cell glass interface. Random media with autocorrelation coefficients representing chromatins are placed into the simulation space. By solving Maxwell equations numerically, the back scattering light intensity field is resolved and used to synthesize simulation PWS images. **(B)** Negative correlation between PWS normalized reflectance and media average RI and phi. n = 10 for each condition.

